# Phenotypic Plasticity in Sub-Antarctic Plants: Costs, Environmental Complexity, and the Facilitative Role of Trait Integration

**DOI:** 10.1101/2023.08.31.555735

**Authors:** Françoise Hennion, Bastien Labarrere, Marine Renaudon, Andreas Prinzing

## Abstract

Phenotypic plasticity is one plant response to climate warming, but it may incur costs and be either limited or facilitated by trait integration or simultaneous environmental shifts such as shading. Empirical studies addressing these costs and limitations across populations or maternal lineages, and how they vary with environmental context, remain scarce. We studied three plant species from the sub-Antarctic Kerguelen Islands, a region undergoing rapid warming, growing seeds from multiple source populations under different temperature and light treatments. We measured plasticity, performance, and morphological integration in young plants. Plants from more plastic source populations performed worse, while those from more integrated populations were more plastic, consistent with the facilitation hypothesis. Shading reduced plasticity in response to warming, and combined shading and warming reduced performance more than single treatments— contrasting with recent findings in Mediterranean plants, where multiple stressors did not limit adaptive plasticity. Our results suggest that phenotypic integration may favour rather than limit plasticity. However, plasticity under climate warming may also be constrained by increased shading from competitors, and does not guarantee success given induced performance costs.

## 1 INTRODUCTION

Climate change is rapidly altering environmental conditions, and as sessile organisms, plants must rely on phenotypic plasticity to cope with these shifts (Sultan 2000). Phenotypic plasticity— the ability of a single genotype to express different phenotypes in response to environmental variation—is a fundamental property of plants, enabling them to persist in heterogeneous and changing environments (Bradshaw 1965, Schlichting 1986, Sultan 2000, Schneider, Bäurle et al. 2026). Plasticity is now widely recognized as a critical component of a species’ adaptive potential to environmental change (Ghalambor, McKay et al. 2007, Callahan, Maughan et al. 2008, Hoffmann and Sgro 2011). However, the expression of a phenotype in a given environment is an integrative outcome of local responses across modular traits and their interactions. Thus, plasticity must be analyzed within the framework of trait variation and co-variation to fully understand phenotypic responses and their evolutionary implications (Pigliucci and Preston 2004, Wang and Zhou 2021, Stotz, Salgado-Luarte et al. 2025).

Plasticity is not inherently adaptive. Only plasticity that produces phenotypes aligned with the direction of natural selection—thereby increasing plant fitness—is considered adaptive (Bell and Galloway 2008). This distinction has gained increasing attention in recent years (Murren, Auld et al. 2015, Molina-Montenegro, Galleguillos et al. 2016, Fox, Donelson et al. 2019, Diamond and Martin 2021). If plasticity is too costly or constrained by other factors, it may reduce rather than enhance performance and fitness (Pigliucci 2005, Auld, Agrawal et al. 2009, Snell-Rood, Kobiela et al. 2018, Wei, Chen et al. 2021). Despite its importance, our understanding of the costs and limitations of plasticity remains incomplete, though recent work has begun to shed light on these trade-offs (Mead, Zavala-Paez et al. 2026).

One major constraint on plasticity is energy allocation. Plastic responses depend on physiological and morphological machinery, including the acquisition of environmental information, signal transduction, and the expression of phenotypic changes (DeWitt, Sih et al. 1998, Vinocur and Altman 2005). These processes incur costs, which can be classified as constitutive (e.g., maintaining sensory and signaling machinery, regardless of whether a plastic response is expressed) or induced (e.g., the energetic and developmental costs of producing new phenotypes, which scale with the intensity of the response) (Callahan, Maughan et al. 2008, Murren, Auld et al. 2015). Such costs can reduce plant fitness and even threaten survival (Sultan and Spencer 2002, Auld, Agrawal et al. 2009). Given the pivotal role of plasticity in plant responses to global change, evaluating plasticity and performance together remains a key challenge (Givnish 2002, Valladares, Gianoli et al. 2007, Auld, Agrawal et al. 2009, Bongers, Olmo et al. 2017).

Another critical limitation arises from the multifactorial nature of environmental variation in natural systems (Valladares, Gianoli et al. 2007). Plastic responses to complex environmental changes are far more difficult to decipher than those studied under controlled laboratory conditions, where typically only one variable is manipulated at a time (Valladares, Gianoli et al. 2007, Callahan, Maughan et al. 2008, Westneat, Potts et al. 2019). In nature, different environmental stressors—such as drought and shading—may elicit opposing plant responses (Sack and Grubb 2002), thereby masking the overall phenotypic adjustment. Moreover, the costs of responding to one stressor may limit responses to others due to energy allocation constraints (Auld, Agrawal et al. 2009). How plant phenotypes adjust to synchronous changes in multiple environmental conditions, and the consequences for performance, remain open questions (Westneat, Potts et al. 2019, Ramos-Muñoz, Blanco-Sánchez et al. 2025).

A further potential constraint on plasticity lies in phenotypic integration—the pattern and magnitude of trait correlations arising from genetic, developmental, and/or functional connections that maintain organismal consistency (Pigliucci and Preston 2004). Environmental change can strongly influence integration (Pigliucci, Whitton et al. 1995, Liu, Schieving et al. 2007, Wood and Brodie 2015, Benavides, Carvalho et al. 2021), and integration has long been assumed to constrain plasticity by limiting the range of variation in traits linked to others (Valladares, Gianoli et al. 2007). However, this relationship is far from resolved (Matesanz, Blanco-Sanchez et al. 2021, Wang and Zhou 2021, Stotz, Salgado-Luarte et al. 2025). Here, we propose an alternative hypothesis: in poorly integrated organisms, extreme plastic modification of one trait may be inconsistent with modifications in other traits, thereby limiting overall plasticity. Conversely, in highly integrated organisms, extreme modification of one trait may align with extreme modifications in others, facilitating plasticity in the first trait and potentially increasing plasticity across traits. Thus, phenotypic integration might both limit and facilitate plasticity, depending on context.

To date, only a few studies have examined how integration affects plasticity. For example, Gianoli and Palacio-Lopez (2009) found that plasticity in response to shading or drought was lowest in traits that were strongly integrated with other traits, supporting the constraint hypothesis. Conversely, Matesanz et al. (2021) showed that phenotypic plasticity of a given trait in response to drought was positively associated with its phenotypic integration with other traits, supporting the facilitation hypothesis. However, these studies remain correlative, as integration cannot be experimentally manipulated without damaging the organism. Importantly, these few studies tried to identify whether more integrated traits are more or less plastic than less integrated traits. However, plasticity varies also among populations or other intraspecific lineages, and to our knowledge no study has tried to identify whether more integrated lineages respond more or less plastically to the environment than less integrated lineages.

Investigating plasticity and integration in response to environmental variation is particularly relevant in regions experiencing drastic climate change. The sub-Antarctic region, characterized by its cool and windy climate year-round, is undergoing rapid and intense environmental shifts (Le Roux and McGeoch 2008, Favier, Verfaillie et al. 2016, Verfaillie, Charton et al. 2021). The Kerguelen Islands, located in the southern Indian Ocean, exemplify this trend, with a mean annual temperature of 4.6°C and mean wind speed of 9.7 m/s at Port-aux-Français (Verfaillie, Charton et al. 2021). Climate change in this region resulted in a significant drying trend associated <ith a marked shift in the storm track location (Favier, Verfaillie et al. 2016, Verfaillie, Favier et al. 2019), and a large increase in the mean temperature (Favier, Verfaillie et al. 2016, Verfaillie, Charton et al. 2021). Direct impacts are already visible, including more frequent and prolonged summer droughts, a rising snow line, and a drastic reduction of the Cook Ice Cap (Hennion, Frenot et al. 2006, Lebouvier, Laparie et al. 2011, Verfaillie, Favier et al. 2015). Plant species in the Kerguelen Islands show signs of stress, such as leaf wilting or increased shoot necrosis, suggesting high sensitivity to ongoing changes (Chapuis, Frenot et al. 2004, Frenot, Lebouvier et al. 2006, Marchand, Tarayre et al. 2021). Due to the geographical isolation of the Kerguelen Islands, species have limited opportunities for migration to track changing conditions, leaving plasticity and adaptive evolution as primary responses (Hoffmann and Sgro 2011).

We studied three indigenous species from the Kerguelen Islands: two *Ranunculus* species—*R. biternatus* and *R. pseudotrullifolius* (Ranunculaceae)—and the Brassicaceae *Pringlea antiscorbutica*. *Ranunculus* species are known for their high plastic responses, particularly heterophylly, which is considered adaptive plasticity (Wells and Pigliucci 2000). Both *R. biternatus* and *R. pseudotrullifolius* exhibit trait plasticity between field and controlled conditions (Hennion, Fiasson et al. 1994). *Pringlea antiscorbutica* shows decreased root or shoot growth in response to salt or heat stress (Hummel, El Amrani et al. 2004, Hummel, Quemmerais et al. 2004) and demonstrates plasticity in shoot growth in response to cultivation conditions (Hennion, Frenot et al. 2006).

In a phytotron experiment, we subjected plants from multiple lineages (populations or maternal lines) of these three species to variations in abiotic conditions, including warming treatments and— for *P. antiscorbutica*—shading treatments and their combination. We measured variation in morphological traits and assessed morphological or physiological performance. Our study aimed to test the following hypotheses regarding the costs, limitations, and integrative constraints of plasticity:

1. Intraspecific lineages with higher plasticity exhibit lower performance under environmental variation (Mead, Zavala-Paez et al. 2026).
2. Intraspecific lineages show lower plasticity in response to two simultaneous environmental changes (e.g., warming + shading) compared to a single condition (Ramos-Muñoz, Blanco-Sánchez et al. 2025).
3. Intraspecific lineages with higher phenotypic integration prior to environmental treatment respond either less plastically or more plastically to that treatment (Matesanz, Blanco-Sanchez et al. 2021, Stotz, Salgado-Luarte et al. 2025).

## 2 MATERIALS AND METHODS

### 2.1 Species under study

The Iles Kerguelen (49°20’00” S, 69°20’00” E) are situated in the southern Indian Ocean within the sub-Antarctic region (Lebouvier and Frenot 2007) (Figure 1). We studied *R. biternatus* Smith, *R. pseudotrullifolius* Skottsberg, and *Pringlea antiscorbutica*. R.Br. ex Hook.f. All are perennial plants. *R. biternatus* and *R. pseudotrullifolius* are two species of austral distribution with a magellanic origin (Lehnebach, Winkworth et al. 2017) while *P. antiscorbutica* is endemic from the South Indian Ocean province (Bartish, Ainouche et al. 2012). On the Iles Kerguelen these species occupy different habitats (Hennion and Walton 1997a). *Ranunculus biternatus* is widespread on the island occurring in habitats below 300m above sea level. In contrast, *R. pseudotrullifolius* being halophilous has more restricted distribution and occurs within a short distance of the coast, occupying peaty or sandy shores and ponds (Hennion and Walton 1997a). On the Iles Kerguelen *P. antiscorbutica* occurs in habitats ranging from coastal meadows to montane fellfields (Hennion and Bouchereau 1998).

**Figure 1.**
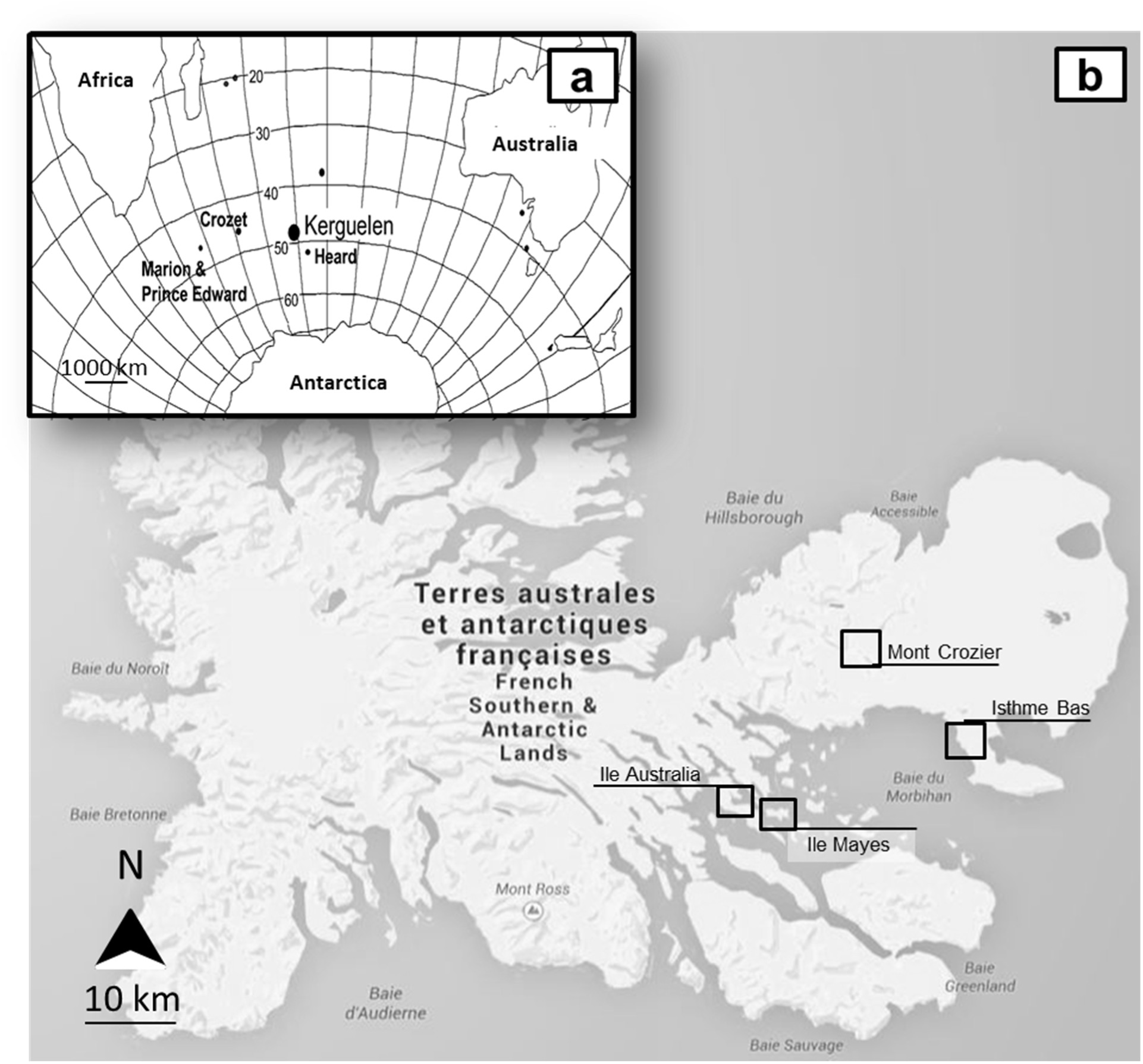
a. Location of Iles Kerguelen in the southern Indian Ocean; b. sampled sites in Iles Kerguelen (map modified from Google maps).

### 2.2 Seed collection

To grow plants under controlled conditions, we used seeds from different natural populations in the three species. Populations were defined as continuous groups of plants living at a given site. Seeds from *P. antiscorbutica* were collected in the field in 2013 in six populations spread across the region covering Ile Australia, Mont Crozier and Ile Mayes (Figure 1). Ten individuals were randomly selected within each population. In each of the ten individuals, fifteen siliques were then randomly collected from the median part of the longest infructescence where seed size is maximum and least variable (Hennion, Schermann-Legionnet and Atlan, unpubl.). Seeds were also collected from *P. antiscorbutica* plants growing from four populations in a phytotron in 2013. In *R. biternatus* and *R. pseudotrullifolius*, seeds were collected in 2014 from 20 individuals across three populations in Isthme Bas, the same in which phenotypic measurements had been performed in 2012. All seeds were stored dry with silica gel at 4°C until their use according to an established protocol (Hennion and Walton 1997b).

### 2.3 Controlled conditions

Seeds were sterilized and germinated in Petri dishes at 25°C for *P. antiscorbutica* and 21°C for *Ranunculus* species, under low light (1kLux) following an established protocol (Hennion and Walton 1997b). To render the overall design more homogeneous we kept a constant number of seedlings from each population. In each species, three hundred seedlings were planted in vermiculite substrate and fertilized with ½ Hoagland solution (Hoagland and Arnon 1938). For the *R. pseudotrullifolius*, NaCl was added to Hoagland solution to reach a concentration of 1g.L^-1^, being consistent with the salinity in the field (Labarrere 2017). Plants were grown hydroponically in a phytotron under conditions (light exposure, photoperiod, temperature and air humidity) mimicking austral summer in Iles Kerguelen at best, according to a previous protocol (Hennion and Bouchereau 1998, Hennion, Frenot et al. 2006). Specifically, plants were grown in a photoperiod of 14h with an irradiance of 15klux from a combination of Philips TLD 79 fluorescent and Philips Aquarelle TLD 89 tubes. Temperature was of 9°C during the day and 4°C during the night with a relative humidity of 75/85% day/night.

### 2.4 Experimental treatments

The experimental design is summarized in Figure 2. Plants were grown under standard, “control” conditions for 5 months at 9°C; 15klux. We then subjected subsets of plants to different treatments during 3 months. In *P. antiscorbutica,* we subjected plants to four conditions: (i) warming (13°C / +4°C), (ii) shading (5klux), (iii) both warming and shading (13°C; 5klux) and (iv) control (9°C; 15klux). In *R. biternatus* and *R. pseudotrullifolius*, we subjected plants to warming (13°C). Due to a low number of available seedlings, *R. biternatus* and *R. pseudotrullifolius* were not subject to shading treatment or combined warming-shading treatment. In *P. antiscorbutica,* a total 320 plants (32 plants from each of 10 source populations) were used: per each of four treatments 8 plants from 10 source populations. In *R. biternatus,* a total 152 plants (8 plants from each of 19 mothers) were grown: per each of two treatments (warming treatment and control) 4 plants from each of 19 mothers for a total of 76 plants. In *R. pseudotrullifolius,* a total 112 plants (8 plants from each of 14 mothers) were grown: per each of two treatments (warming treatment and control) 4 plants from 14 mothers for a total of 56 plants. Placement of plants within a treatment was random with respect to levels of integration or of performance, thereby avoiding any confounding effect of possible within-treatment variation of microenvironmental conditions and how different lineages of plants responded to a given treatment. Plants were all 8-months old at the end of experiments.

**Figure 2.**
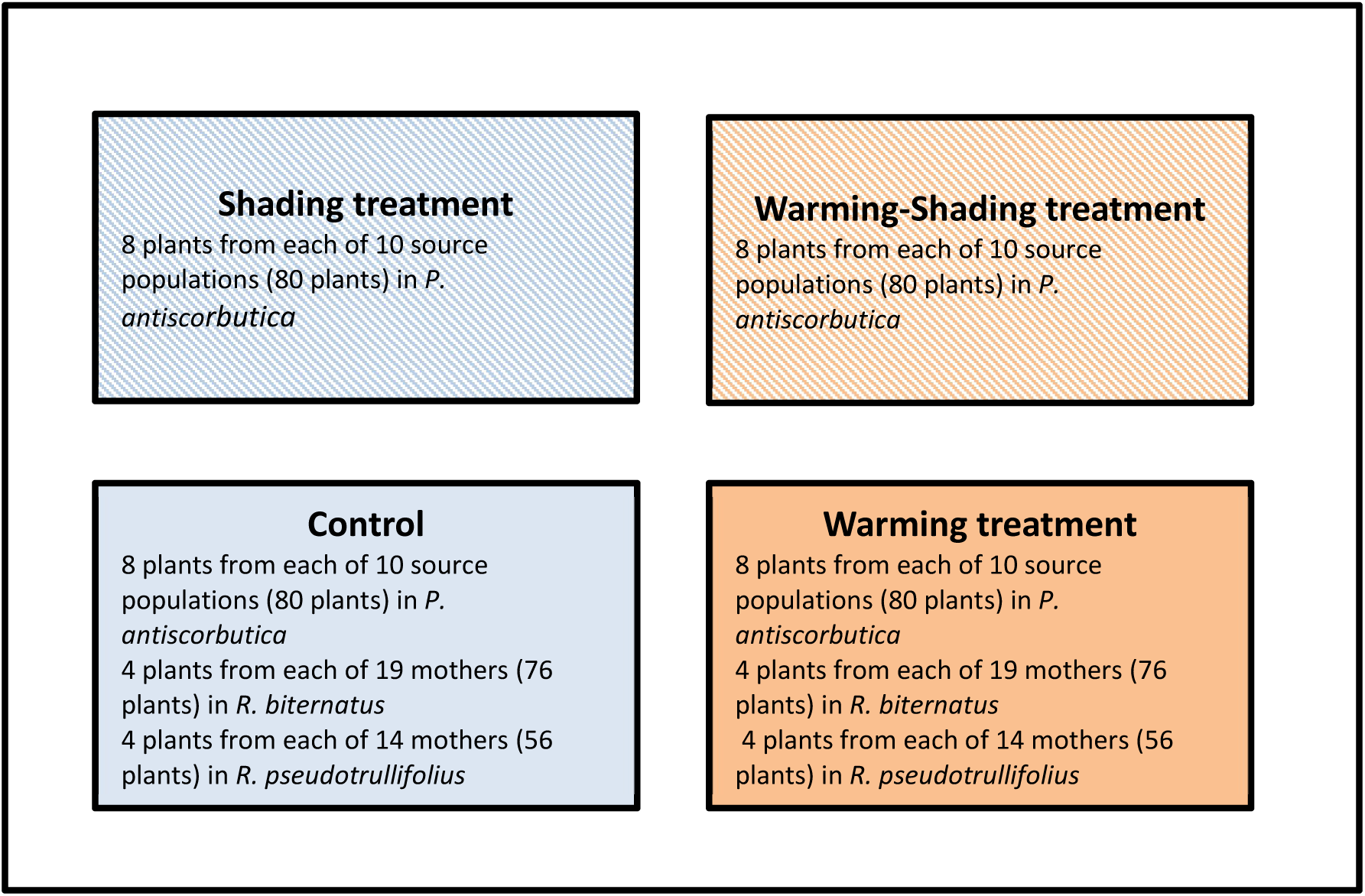
Illustration of experimental design. The numbers of plants used per treatment, per species are shown.

### 2.5 Trait measurements

We conducted two sets of plant measurements. First, we measured plants after 5 months growth while all plants were subjected to control conditions to determine baseline differences between plants (T0). We measured plants again after 3 months growth in the different abiotic conditions (T1). Traits considered were height, diameter and number of leaves of the rosette, and of the 2 longest leaves: length and width of lamina and length of petiole. These traits were shown to respond to increased temperature in previous work (Hennion, Frenot et al. 2006, Hermant, Prinzing et al. 2013, Labarrere 2017). In *R. biternatus* and *R. pseudotrullifolius*, diameter changed during plant handling and therefore was not measured. We also estimated physiological and morphological plant performances. To determine physiological performance, we used the maximum efficiency of photosystem II (Maxwell and Johnson 2000, Zhu, Long et al. 2008), a useful performance parameter known to decrease with stress (Maxwell and Johnson 2000, Kalaji, Bosa et al. 2011). Maximum PSII efficiency (Fv/Fm) was measured using a PAM chlorophyll fluorometer system (Heinz Walz, Effeltrich, Germany) in saturating pulse mode. After 30 minutes of dark adaptation, the maximum fluorescence of leaves was measured under dark conditions (Fm) and a subsequent saturating pulse of light was applied. Minimum fluorescence (F0) was determined under weak light before a pulse, and variable fluorescence (Fv=Fm–F0) was calculated. To determine morphological performance, we used leaf spread (Zamora, Gómez et al. 1998). Leaves that are not fully spread, but curled, indicated that the plant is not fully performant. Part of leaf that is curled is unexposed to light and therefore does not contribute to photosynthesis. We measured leaf spread as the ratio between length of leaf that is not curled and maximum flattened leaf length.

### 2.6 Data analyses

#### 2.6.1 Trait means and plasticity

We investigated plant plasticity comparing plant trait values between the different experimental conditions, *i.e.* a trait change, with in whatever direction, with a change in environment. Prior to calculating plasticity of traits, we aimed to remove the intrinsic trait value differences among plants to correct for differences between plants already present prior to exposing them to controls or treatments during months 5 – 8. Thus, we calculated trait value as standardized trait value Ts = (T1-T0)/T0. With T0 being the measurement of the trait done during the first set of measurements - when all plants were subjected to the same conditions – and T1 being the measurement of the trait done during the second set of measurements, when plants were subjected to different experimental conditions. In *P. antiscorbutica*, we calculated trait means of these standardized trait values across plants from the same population subjected to same condition. In *R. biternatus* and *R. pseudotrullifolius*, we calculated trait means across plants from the same mother subjected to same condition. So, to summarize, we had one data point per maternal line (or population) and treatment. In each species, we calculated - for each population (or mother) and treatment - trait plasticity as the absolute value of the difference between trait mean under treatment and trait mean under control. In *P. antiscorbutica* we also calculated the mean degree of plasticity across the three treatments. Samples sizes did not permit the same calculus for *Ranunculus* species. These means across individuals per population or per mother could then be compared to phenotypic integration calculated across individuals per population or per mother. For consistency we used source populations or mothers as our level of analyses in all further comparisons.

#### 2.6.2 Phenotypic integration

We calculated phenotypic integration across *P. antiscorbutica* individuals from the same source populations. We did so prior to treatments, thereby avoiding any effect of environment on integration (Wang and Zhou 2021). Samples sizes did not permit the same calculus for *Ranunculus* species. We focused on the dominant axis of phenotypic integration, quantified as the percentage of variance explained by the first axis of a Principal Component Analysis (PCA) performed on traits (Tucic, Vuleta et al. 2013, Haber and Dworkin 2017, Michaud, Veron et al. 2020). PCA was performed using correlation matrix. Traits were height and diameter of the rosette, and of the 2 longest leaves: length and width of lamina and length of petiole, leaf 1 older than leaf 2. Trait values were scaled and PCA conducted using the FactoMineR package of R 4.0 (Lê, Josse et al. 2008, R 2020). Note that all PCAs are calculated across the same number of traits hence the same total number of axes, rendering % explained variance directly comparable among PCAs. We investigated whether degree of phenotypic integration differed before and after the exposure to treatment, but found no differences [Supplementary Table S2a]. Moreover, we found that degree of phenotypic integration did not differ among treatments [Supplementary Table S2b].

#### 2.6.3 Hypothesis Testing

The relationships between trait means, plasticity or integration were tested using simple linear regression. We determined differences of trait values among treatments using ANOVA. We used MANOVA to determine whether plant phenotype (accounting for all traits) differed among treatments. The same traits were tested in MANOVA and ANOVA analyses. Analyses were conducted in R 4.0(R 2020). For multiple testing on the same data set, p-values were corrected using sequential Bonferroni’s correction (Cabin and Mitchell 2000). In all regression analyses we verified the assumptions of the analyses using residual plots.

## 3 RESULTS

### 3.1 Effects of treatments on plant traits

In *P. antiscorbutica*, plant phenotype of maternal lineages was significantly larger in warming treatment than in control (p=0.005; F=5.9; dferror=16; MANOVA). In contrast, plant phenotype was significantly smaller in response to shading or combined warming-shading treatments than in control (respectively: p=0.0043; F=6.8; dferror =16 and p=0.012; F=5.1; dferror =16; MANOVA). We did not find any difference in plant phenotype between shading and combined warming-shading treatments (p=0.55; F=0.93: dferror =16; MANOVA). Differences of individual traits between treatments and the control are indicated Table 1.

**Table 1.**
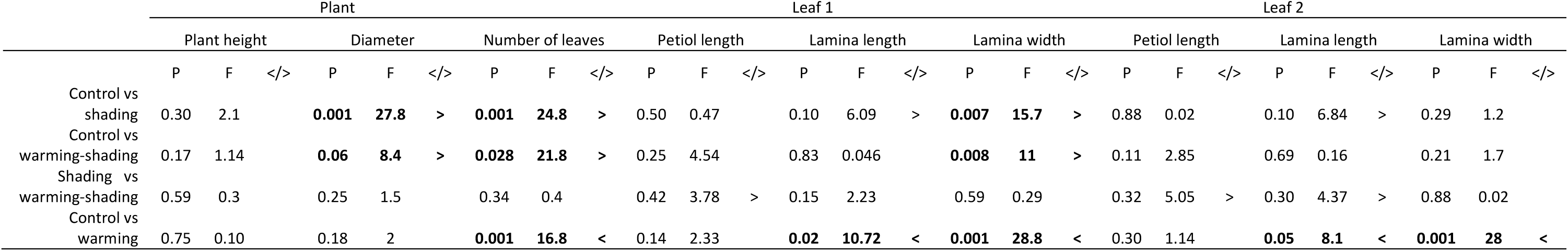
ANOVA analyses indicating trait differences between different abiotic treatments in *P. antiscorbutica*. P-values (P) after sequential Bonferroni’s correction and F values (F) are indicated. "</>": trait is lower/larger in the first treatment indicated. dferror =18 source populations.

|  | Plant |  |  |  |  |  |  |  |  | Leaf 1 |  |  |  |  |  |  |  |  | Leaf 2 |  |  |  |  |  |  |  |  |
|---|---|---|---|---|---|---|---|---|---|---|---|---|---|---|---|---|---|---|---|---|---|---|---|---|---|---|---|
|  | Plant height |  |  | Diameter |  |  | Number of leaves |  |  | Petiol length |  |  | Lamina length |  |  | Lamina width |  |  | Petiol length |  |  | Lamina length |  |  | Lamina width |  |  |
|  | P | F | </> | P | F | </> | P | F | </> | P | F | </> | P | F | </> | P | F | </> | P | F | </> | P | F | </> | P | F | </> |
| Control vs shading | 0.30 | 2.1 |  | **0.001** | **27.8** | **>** | **0.001** | **24.8** | **>** | 0.50 | 0.47 |  | 0.10 | 6.09 | > | **0.007** | **15.7** | **>** | 0.88 | 0.02 |  | 0.10 | 6.84 | > | 0.29 | 1.2 |  |
| Control vs warming-shading | 0.17 | 1.14 |  | **0.06** | **8.4** | **>** | **0.028** | **21.8** | **>** | 0.25 | 4.54 |  | 0.83 | 0.046 |  | **0.008** | **11** | **>** | 0.11 | 2.85 |  | 0.69 | 0.16 |  | 0.21 | 1.7 |  |
| Shading vs warming-shading | 0.59 | 0.3 |  | 0.25 | 1.5 |  | 0.34 | 0.4 |  | 0.42 | 3.78 | > | 0.15 | 2.23 |  | 0.59 | 0.29 |  | 0.32 | 5.05 | > | 0.30 | 4.37 | > | 0.88 | 0.02 |  |
| Control vs warming | 0.75 | 0.10 |  | 0.18 | 2 |  | **0.001** | **16.8** | **<** | 0.14 | 2.33 |  | **0.02** | **10.72** | **<** | **0.001** | **28.8** | **<** | 0.30 | 1.14 |  | **0.05** | **8.1** | **<** | **0.001** | **28** | **<** |

In *R. biternatus*, number of leaves as well as petiole length and lamina width of leaf 2 of populations were significantly larger in warming treatment compared to control (Table 2). In *R. pseudotrullifolius*, lamina length of leaf 1 was significantly larger in warming treatment than in control (Table 2). Overall, phenotype was larger (marginally significant) in warming in *R. biternatus* but not in *R. pseudotrullifolius* (respectively: p=0.058; F=2.2; dferror =34 and p=0.57; F=0.8; dferror =24; MANOVA).

**Table 2.** ANOVA analyses indicating trait differences between warming and control in *R. biternatus* and *R. pseudotrullifolius*. P-values (P) after sequential Bonferoni’s correction and F values (F) are indicated. "</>": trait is lower/larger in the control. In *R. biternatus*, dferror = 36 mothers; in *R. pseudotrullifolius*, dferror =26 mothers.

|  | Plant |  |  |  |  |  | Leaf number 1 |  |  |  |  |  |  |  |  | Leaf number 2 |  |  |  |  |  |  |  |  |
|---|---|---|---|---|---|---|---|---|---|---|---|---|---|---|---|---|---|---|---|---|---|---|---|---|
|  | Plant height |  |  | Number of leaves |  |  | Petiol length |  |  | Lamina length |  |  | Lamina width |  |  | Petiol length |  |  | Lamina length |  |  | Lamina width |  |  |
| Control vs warming | P | F | </> | P | F | </> | P | F | </> | P | F | </> | P | F | </> | P | F | </> | P | F | </> | P | F | </> |
| *R. biternatus* | 0.14 | 3.62 |  | **0.0091** | **10.94** | **<** | 0.085 | 4.12 |  | 0.12 | 2.44 |  | 0.11 | 4.67 |  | **0.03** | **7.19** | **<** | 0.066 | 1.07 |  | **0.0003** | **5.8** | **<** |
| *R. pseudotrullifolius* | 0.54 | 0.66 |  | 1 | 1.36 |  | 0.43 | 0.18 |  | **0.033** | **3.1** | **<** | 0.077 | 2.6 |  | 0.85 | 1 |  | 0.70 | 0.39 |  | 0.90 | 0.6 |  |

### 3.2 Effects of treatments on plant performance

In *P. antiscorbutica*, we found that morphological performance (percentage of leaf spread) of maternal lineages was lower in combined warming-shading treatment than in the other treatments or in the control (Table 3). In *R. biternatus* we found that plant performances of populations did not differ significantly between control and warming treatment (Table 4). In *R. pseudotrullifolius,* we found that physiological performance of populations was higher in warming treatment. Yet, this result is only marginally significant after sequential Bonferroni’s correction (Table 4).

**Table 3.** ANOVA analyses indicating differences of performance (photosynthetic efficiency and leaf spread) between treatments in *P. antiscorbutica*. P-values (P) after sequential Bonferroni’s correction and F values (F) are indicated. "</>": performance is lower/higher in the first treatment indicated. dferror =18 source populations.

|  | Photosynthetic efficiency |  |  | Leaf spread |  |  |
|---|---|---|---|---|---|---|
|  | P | F | </> | P | F | </> |
| Control vs warming | 0.11 | 2.8 |  | 0.90 | 0.014 |  |
| Control vs shading | 0.19 | 1.81 |  | 0.09 | 3.32 |  |
| Control vs warming-shading | 0.24 | 1.51 |  | **0.0002** | **31.74** | **>** |
| Shading vs warming-shading | 0.58 | 0.32 |  | **0.02** | **8.02** | **>** |
| Warming vs warming-shading | 0.42 | 0.69 |  | **0.0002** | **27.92** | **>** |

**Table 4.** ANOVA analyses indicating differences of performance (percentage of survival, photosynthetic efficiency and leaf spread) between warming and control in *R. biternatus* and *R. pseudotrullifolius*. P-values (P) after sequential Bonferroni’s correction and F values (F) are indicated. "</>": performance is lower/higher in the control. In *R. biternatus*, dferror = 36 mothers; in *R. pseudotrullifolius*, dferror =26 mothers.

|  |  | Percentage of survival |  |  | Photosynthetic efficiency |  |  | Leaf spread |  |  |
|---|---|---|---|---|---|---|---|---|---|---|
|  |  | P | F | </> | P | F | </> | P | F | </> |
| *R. biternatus* | Control vs warming | 0.14 | 1.01 |  | 0.90 | 0.014 |  | 0.51 | 0.2 |  |
| *R. pseudotrullifolius* | Control vs warming | 0.14 | 0.93 |  | 0,08 | 3.24 | < | 0.11 | 1.65 |  |

### 3.3 Effect of treatments on the degree of plasticity

In *P. antiscorbutica*, we found that the degree of plasticity of maternal lineages was higher in response to warming treatment than in response to shading treatment or combined warming-shading treatment (respectively: p=0.05; F=2.19; dferror =16 and p=0.036; F=2.21; dferror =16; MANOVA). In contrast, the degree of plasticity did not differ between shading treatment and combined warming-shading treatment (p=0.45; F=0.46; dferror =16; MANOVA). Specifically, degrees of plasticity of plant height and lamina width of the tallest leaf were higher in response to warming treatment than to shading treatment (Table 5). Degrees of plasticity of lamina width of leaf 1 and leaf 2 were higher in response to warming treatment than in response to combined warming-shading treatments (Table 5).

**Table 5.**
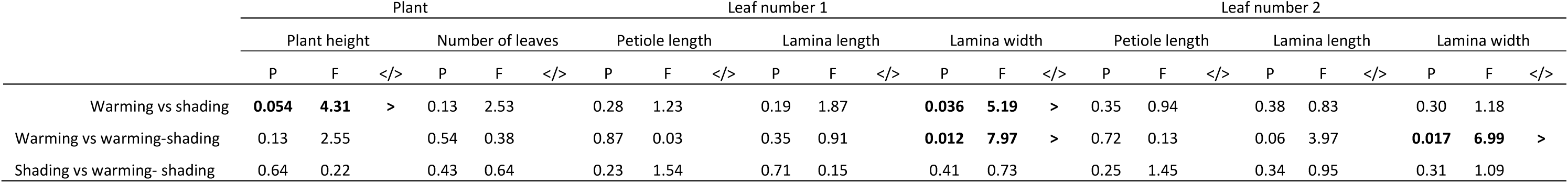
ANOVA analyses indicating differences of the degree of plasticity between treatments in *P. antiscorbutica*. "</>": trait is less/more plastic in the first treatment indicated. P-values (P) and F values (F) are indicated. dferror =16 source populations.

### 3.4 Relationship between degree of plasticity and performance

In *P. antiscorbutica*, we found significant negative relationship between the degree of trait plasticity of maternal lineages and plant performance (percentage of leaf spread and marginally photosynthetic efficiency) in response to warming treatment (Figure 3). Also, we found a significant negative relationship between trait plasticity and performance (percentage of leaf spread) in response to combined warming-shading treatment (Figure 4). We did not find any significant relationships between the degree of trait plasticity and plant performances in response to shading treatment (p-value>0.05; all adj.r²<0.1; simple linear regressions). To explore constituent costs of being plastic we compared across-treatment means of plasticity per source population to performances in control condition, where no plasticity is expressed. We found no relationships (p-value>0.05; all adj.r²<0.1; simple linear regression).

**Figure 3.**
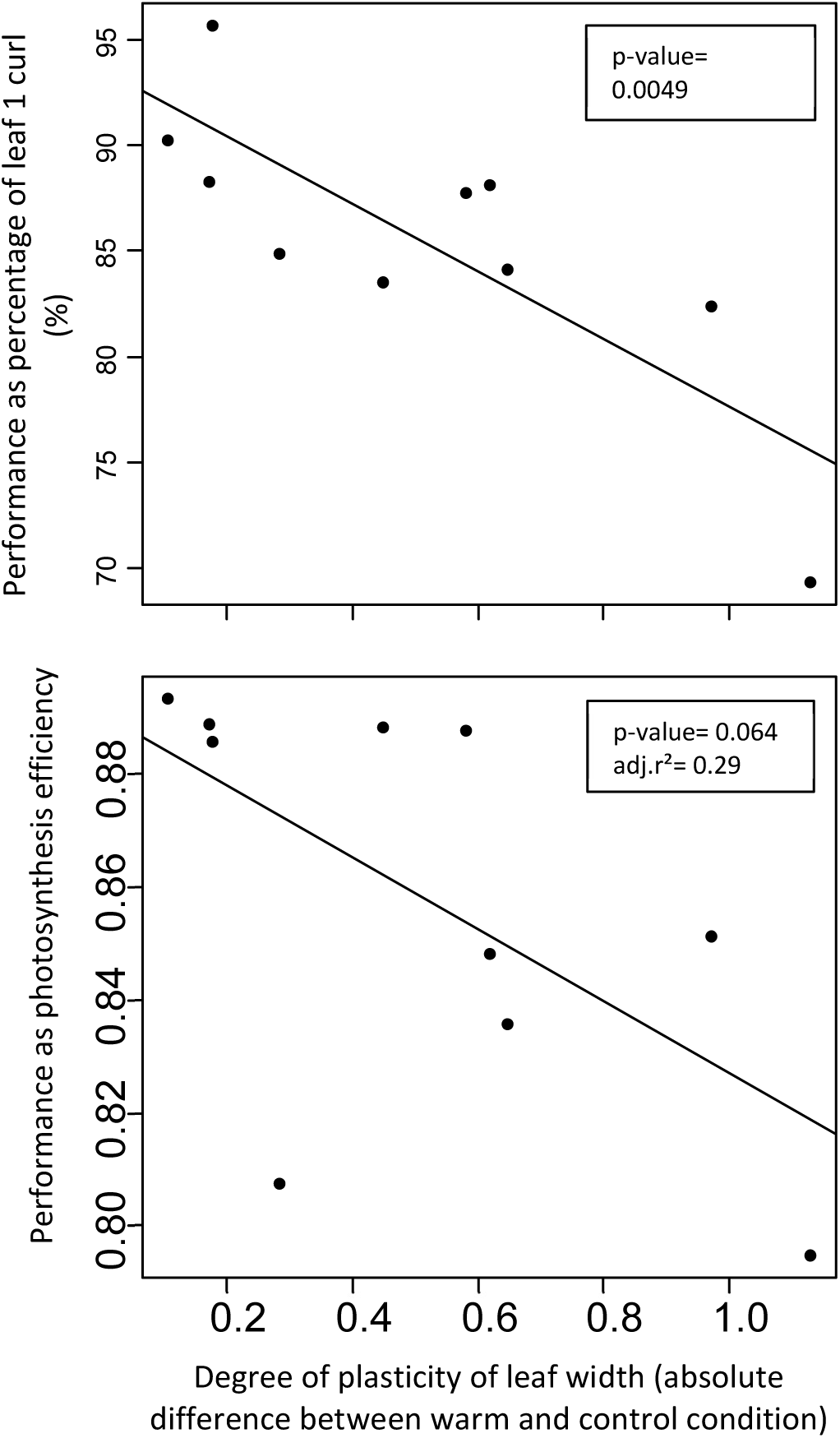
Negative relationship between the degree of plasticity and performance in the warming treatment across maternal lineages of *P. antiscorbutica*. P-values and adj.r² are indicated, N=9.

**Figure 4.**
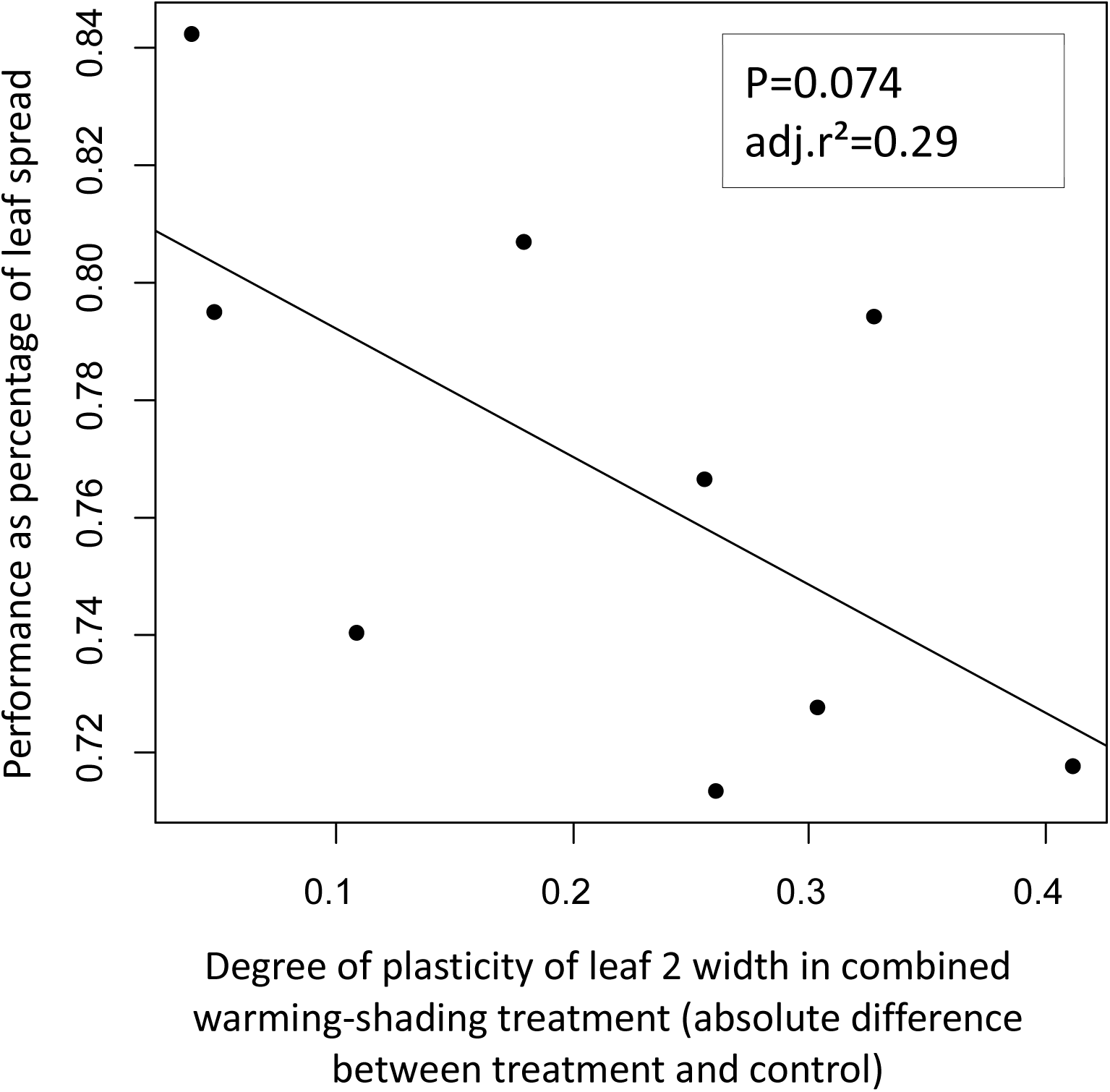
Negative relationship between the degree of plasticity and performance in the warming-shading treatment across maternal lineages of *P. antiscorbutica*. P-values and adj.r² are indicated, N=9.

In *R. biternatus* and *R. pseudotrullifolius*, we did not find any significant relationship between the degree of trait plasticity of populations and plant performance.

### 3.5 Relationship between phenotypic integration prior to treatment and the degree of plasticity in response to treatment

In *P. antiscorbutica*, we found a significant positive relationship between phenotypic integration of maternal lineages and the degree of plasticity of leaf 1 length in shading treatment (Figure 5). Also, we found marginally significant positive relationships between phenotypic integration and the degree of plasticity of leaf 2 length in shading treatment (Figure 5). We found no significant relationship between integration and the degree of trait plasticity in other treatments.

**Figure 5.**
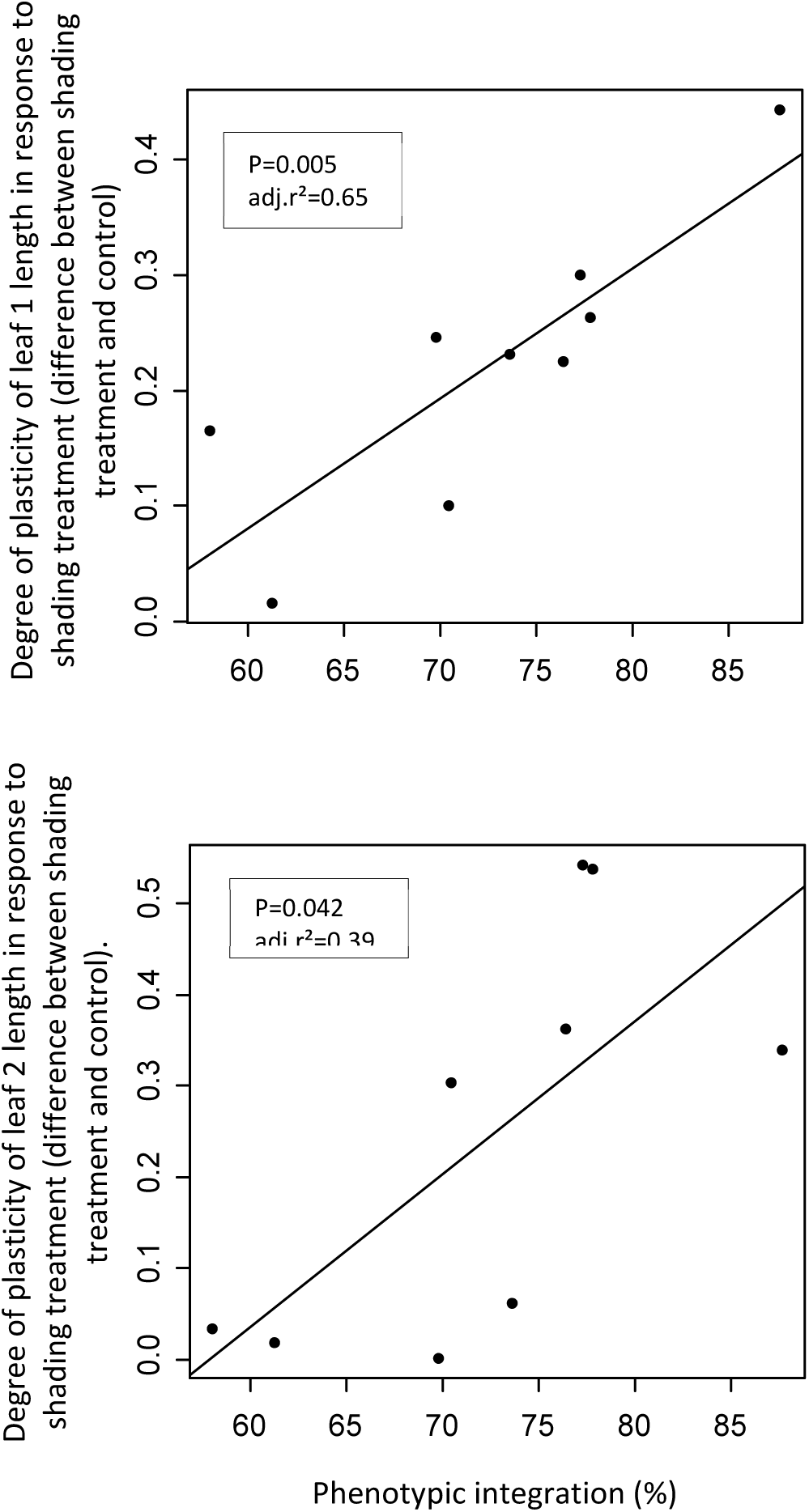
Positive relationship between phenotypic integration and the degree of plasticity of leaf 1 width or leaf 2 length in the shading treatment across maternal lineages of *P. antiscorbutica*. N=9.

## 4 DISCUSSION

As sessile organisms, plants must cope with changing environmental conditions, and phenotypic plasticity is widely recognized as a primary mechanism for such response (Ghalambor, McKay et al. 2007, Fox, Donelson et al. 2019). However, while plasticity enables rapid responses to environmental variation, it is not without costs or limitations. Theoretical frameworks have long acknowledged that plasticity may incur fitness costs (Pigliucci 2005, Valladares, Gianoli et al. 2007), yet empirical evidence for these trade-offs remains limited (Pigliucci 2005, Valladares, Gianoli et al. 2007, Callahan, Maughan et al. 2008, Murren, Auld et al. 2015, Molina-Montenegro, Galleguillos et al. 2016, Wei, Chen et al. 2021). Furthermore, most studies to date have focused on the independent responses of individual traits to single environmental factors (Valladares, Gianoli et al. 2007, Matesanz, Blanco-Sanchez et al. 2021), despite the fact that plants in nature face multifactorial environmental variation and exhibit phenotypic integration—where traits are developmentally, genetically, or functionally correlated and cannot vary independently (Pigliucci and Preston 2004, Wang and Zhou 2021).

In this study, we investigated the relationships between plasticity, performance, and phenotypic integration under controlled environmental treatments. Our results reveal three key insights: (i) plasticity can decrease performance, suggesting strong induced costs; (ii) plasticity is reduced when plants respond to combined abiotic conditions compared to single-factor variations; and (iii) phenotypic integration can facilitate plasticity in certain contexts, particularly under shading. These findings contribute to a growing body of literature that highlights the context-dependence of plasticity and its complex interplay with trait integration (Stotz, Salgado-Luarte et al. 2025, Schneider, Bäurle et al. 2026).

### 4.1 Trait Values Under Different Treatments

The Kerguelen Islands are experiencing rapid and intense climate change, characterized by rising temperatures and reduced rainfall (Lebouvier, Laparie et al. 2011, Favier, Verfaillie et al. 2016, Verfaillie, Charton et al. 2021). Native plants in this region already exhibit stress symptoms, such as leaf wilting and increased shoot necrosis during dry summer periods (Chapuis, Frenot et al. 2004, Frenot, Lebouvier et al. 2006, Marchand, Tarayre et al. 2021). Previous work has documented plastic responses in the species studied here (Hennion, Fiasson et al. 1994, Hennion and Walton 1997a, Hummel, Quemmerais et al. 2004). In our experiment, all three species exhibited plastic morphological responses to abiotic changes. In *Pringlea antiscorbutica*, four out of nine traits increased in response to warming, consistent with earlier observations at the seedling stage, where shoot growth increased under warm temperature treatments (22°C night/ 25°C day) compared to cooler conditions mimicking Kerguelen summer (5°C night/ 10°C day) (Hummel, El Amrani et al. 2004). Similarly, *Ranunculus biternatus* showed increased values for three out of eight traits under warming, while *R. pseudotrullifolius* exhibited plasticity in only one out of eight traits, which increased in response to warming. These species-specific differences in plastic responses underscore the diverse strategies employed by sub-Antarctic flora to cope with environmental variation.

### 4.2 Degrees of Plasticity Under Simultaneous Environmental Change

A major challenge in plasticity research is understanding how plants respond to multifactorial environmental variation (Westneat, Potts et al. 2019). In *P. antiscorbutica*, we investigated how the degree of plasticity varied across treatments, particularly whether simultaneous variation in multiple environmental conditions (e.g., warming + shading) would limit plasticity compared to single-factor changes. We found that plasticity was higher in response to warming than to shading, though we acknowledge that the two treatments differ in units and intensity, making direct comparisons challenging. More importantly, plasticity was lower in response to the combined warming-shading treatment than to warming alone.

One possible explanation for this reduced plasticity is that warming and shading have opposite effects on plant growth, potentially canceling each other out and resulting in a lower overall plastic response (Sack and Grubb 2002). Alternatively, resource allocation constraints may prevent traits from responding fully to multiple environmental cues (Auld, Agrawal et al. 2009). Interestingly, we observed that plant phenotypes under the combined warming-shading treatment were closer to those under shading alone than to warming alone, suggesting that shading imposes a stronger constraint than warming. This aligns with the emerging view that multidimensional plasticity cannot be fully understood by extrapolating from unidimensional models (Westneat, Potts et al. 2019).

However, our findings contrast with recent work on Mediterranean plants, where Ramos-Muñoz, Blanco-Sánchez et al. (2025) demonstrated that warming and competition (which presumably involves shading) did not limit adaptive plastic responses to drought. This discrepancy highlights the context-dependence of plasticity, which may vary with species, environmental conditions, or the nature of the stressors involved. In sub-Antarctic ecosystems like Kerguelen, where plants are already operating at the edge of their physiological tolerances (Chapuis, Frenot et al. 2004, Frenot, Lebouvier et al. 2006, Marchand, Tarayre et al. 2021), the combined effects of warming and shading may exceed the plastic capacity of native species, whereas Mediterranean plants may possess greater inherent plasticity to multiple stressors.

### 4.3 Plasticity in the Face of Climate-change-induced Warming and Shading by Competitors

Species that exhibit plasticity in response to abiotic conditions are often considered better equipped to face climate change (Valladares, Gianoli et al. 2007). In the Kerguelen Islands, mean temperatures are rising (Favier, Verfaillie et al. 2016, Verfaillie, Charton et al. 2021), and the plasticity observed in *P. antiscorbutica* and *R. biternatus* in response to warming may enhance their capacity to cope with these changes. In contrast, *R. pseudotrullifolius* exhibited limited plasticity in response to temperature, suggesting it may be less resilient to warming. Notably, in all three species, plasticity did not decrease performance over the short term (three months). However, we acknowledge that long-term exposure to elevated temperatures may eventually reduce performance, as suggested by previous experiments. For instance, under well-watered cultivation in a common garden with a temperate climate (warmer than sub-Antarctic conditions) in Brest, western Brittany, France, young plants of *P. antiscorbutica*, *R. biternatus*, and *R. pseudotrullifolius* initially showed increased growth but ultimately collapsed after several months (Hennion, pers. comm.).

In the field, plants are also exposed to additional stressors such as wind, variable soil water saturation, and nutrient limitations. Combined with these factors, increased temperatures may further impact performance. Rising temperatures in the Kerguelen Islands are also expected to favor the expansion of invasive plant species (Frenot, Lebouvier et al. 2006), which may compete with native species for resources such as light. In *P. antiscorbutica*, we found that phenotypes under shading alone were similar to those under combined warming and shading, with individual traits decreasing in both cases. While neither shading nor warming alone affected performance, the combination of warming and shading significantly reduced performance. This suggests that *P. antiscorbutica* may be particularly sensitive to the interaction between climate change and light competition, whether from native or introduced species. Importantly, warming itself may trigger increased plant growth, which could exacerbate light competition and further constrain plasticity (Ramos-Muñoz, Blanco-Sánchez et al. 2025). Our results thus highlight a potential vicious cycle: warming promotes growth, which increases shading, which in turn limits the ability of plants to respond plastically to further environmental changes.

### 4.4 Phenotypic Integration and Plasticity

The relationship between phenotypic integration and plasticity remains an open question in evolutionary ecology (Matesanz, Blanco-Sanchez et al. 2021, Wang and Zhou 2021). Before assessing how integration influences plasticity, it is essential to determine whether integration itself responds to environmental conditions. In our study, we found no response of integration to abiotic treatments, suggesting that integration may instead modulate plastic responses to these conditions.

Traditionally, integration has been assumed to constrain plasticity by limiting the range of variation in traits that are developmentally or functionally linked (Gianoli and Palacio-López 2009). However, we proposed an alternative hypothesis: in highly integrated organisms, extreme modifications in one trait may align with extreme modifications in other traits, thereby facilitating overall plasticity while maintaining organismal consistency. Our results provide partial support for this hypothesis. We found no relationship between pre-treatment integration and plasticity in response to warming or combined warming-shading treatments. However, maternal lineages with higher integration prior to treatment exhibited greater plasticity in response to shading alone. This suggests that integration can facilitate plastic responses to certain environmental changes, particularly those that impose strong constraints (e.g., shading).

Our findings align with recent meta-analytical evidence showing that the relationship between integration and plasticity is highly context-dependent. While many studies report a negative association (integration constraining plasticity), others—including ours—find positive or neutral relationships (Stotz, Salgado-Luarte et al. 2025). For example, Matesanz, Blanco-Sánchez et al. (2021) showed that plasticity in response to drought was positively associated with trait integration, supporting the facilitation hypothesis. To our knowledge, our study is the first to demonstrate that lineages with higher integration respond more plastically to environmental change, suggesting that plastic changes in one trait may be easier when they are integrated with other traits. However, we caution that our study is correlative, and we cannot rule out the possibility that both integration and plasticity are influenced by a third, unknown factor. Future work using experimental manipulations of integration (e.g., through genetic or developmental approaches) would be valuable to disentangle these relationships.

### 4.5 Costs of Plasticity: High Plasticity Relates to Decreased Performance

While plasticity enables plants to cope with environmental changes, it is not without costs, which can compromise fitness (Valladares, Gianoli et al. 2007, Bell and Galloway 2008, Auld, Agrawal et al. 2009, Murren, Auld et al. 2015). We investigated whether the degree of plasticity in populations or maternal lineages was associated with increased or decreased performance in *P. antiscorbutica*. We found that plasticity statistically decreased performance under high-temperature treatment, while no significant relationship was observed in other treatments. Overall, our results provide empirical evidence for the costs of plasticity, and these costs appear to be environment-dependent.

Plasticity costs can be broadly categorized into constitutive and induced costs (Sultan and Spencer 2002, Callahan, Maughan et al. 2008). Constitutive costs arise from maintaining the physiological machinery required for plasticity, even in stable environments, while induced costs result from the energetic and developmental expenses of producing new phenotypes in response to environmental change (Auld, Agrawal et al. 2009). In our study, performance under stable conditions was not related to the capacity for plastic responses, suggesting that constitutive costs may be minimal in our system. In contrast, induced costs—reflecting the resources required for morphological modifications—appear to play a significant role. We found that higher plasticity was negatively correlated with performance post-treatment, particularly under warming. Given that warming induced a higher degree of plasticity than other treatments, we suggest that high plasticity generates substantial induced costs, which in turn reduce performance.

We also observed a negative plasticity-performance relationship under the combined warming-shading treatment. This aligns with the idea that more stressful environments, such as those combining multiple stressors, reduce the energy available for growth and plasticity (Auld, Agrawal et al. 2009). Critically, the complete absence of a positive relationship between plasticity and performance in our system suggests that more plastic individuals do not necessarily perform better under environmental change. On the contrary, highly plastic individuals may perform poorly, with costs manifesting primarily as induced costs once the environmental change occurs. These findings are consistent with recent work by Mead et al. (2026), who demonstrated that plasticity evolved for climate variability can constrain performance under warming, and with broader reviews emphasizing that plasticity alone is insufficient for long-term adaptation (Schneider, Bäurle et al. 2026).

## 5 CONCLUSIONS

Despite decades of research, our understanding of the costs and limits of plasticity in plants remains incomplete (Pigliucci 2005, Murren, Auld et al. 2015, Matesanz, Blanco-Sanchez et al. 2021). In this study, we provide empirical evidence that simultaneous shifts in multiple environmental factors can limit plasticity in individuals from different lineages, potentially reducing the capacity of populations to respond to climate warming, particularly under increased light competition. Additionally, we found evidence for a positive relationship between phenotypic plasticity and pre-treatment phenotypic integration in response to shading, suggesting that trait changes may be more plastic when they are well-integrated into the organism’s body plan. Finally, we demonstrated that highly plastic individuals often perform poorly, likely due to induced costs of plasticity.

Our results highlight that plastic responses do not guarantee success in new environments. In some cases, less plastic individuals may perform as well as, or even better than, highly plastic individuals under environmental change. This underscores the need for future studies to explore whether the relationships we observed hold for traits beyond vegetative morphology, as well as for other environmental stressors. Moreover, given the context-dependence of plasticity and integration (Ramos-Muñoz, Blanco-Sánchez et al. 2025, Stotz, Salgado-Luarte et al. 2025), research should prioritize experimental manipulations of integration and long-term studies to better understand the mechanisms underlying these relationships. In sub-Antarctic ecosystems like Kerguelen, where plants face rapid climate change and limited migration opportunities, such insights are critical for predicting species resilience and guiding conservation efforts.

## Supplementary Materials

Supplementary information is available alongside the main manuscript and consist of the following. Table S1: MANOVA analyses indicating trait differences among source mothers (*R. biternatus* and *R. pseudotrullifolius*) or source populations (*P. antiscorbutica*) in the control condition. Table S2: ANOVA analyses indicating in *P. antiscorbutica* whether a) degree of phenotypic integration differs before and after application of treatments, b) phenotypic integration differs among treatments.

## Author Contributions

FH, BL and AP conceived the ideas. FH and BL performed the fieldwork. BL and MR performed the laboratory work. BL, MR and AP analysed the data. BL, FH and AP led the writing. All authors contributed to discussions.

## Funding

Field work was supported by IPEV (programs 1116 PlantEvol - F. Hennion and 136 Ecobio - M. Lebouvier). Laboratory work was supported by CNRS Zone-Atelier Antarctique, CNRS INEE grant (PICS “AntarctBiodiv” to FH) and University Rennes 1 (Direction de la Recherche et de l’Innovation). B.L. was supported by a PhD grant from Ministry of Research and Education (France).

## Data Availability Statement

The datasets generated during the current study will be available online in Zone-Atelier Antarctique (ZATA) repository in data.InDoRES.

## Acknowledgments

We thank IPEV programs 136 Ecobio (M. Lebouvier), 1116 PlantEvol (F. Hennion) and Réserve Naturelle TAF for contribution to seed collection in Iles Kerguelen. We thank personnel of common facility ECOLEX, ECOBIO Rennes (Valérie Gouesbet, Jean-Luc Foulon, Thierry Fontaine-Breton and †Fouad Nassur) for help in plant cultivation. This research is linked to CNRS Zone-Atelier Antarctique (ZATA). During the preparation of this manuscript, the author(s) used Vibe (Mistral AI), for the purposes of updating the bibliography, refining the narrative to align with recent literature, and improving clarity, conciseness and flow. The authors have reviewed and edited the output and take full responsibility for the content of this publication.

## Conflict of interest statement

The authors declare no conflict of interest.

## Notes

### Competing Interest Statement

The authors have declared no competing interest.

### Summary of Updates

Bibliography was updated and the narrative was refined to align with the most recent literature and improve clarity and flow.

