## Supplemental Tables 1 and 2 for "Phenotypic Plasticity in Sub-Antarctic Plants: Costs, Environmental Complexity, and the Facilitative Role of Trait Integration"

Table S1: MANOVA analyses indicating trait differences among source mothers (*R. biternatus* and *R. pseudotrullifolius*) or source populations (*P. antiscorbutica*) in the control condition. Traits used in the analyses are: plant height, number of leaves, and in the 2 longest leaves: petiole length, lamina length and lamina width. P-values (P), F values (F) and error degree of freedom (dferror) indicated.

| | P | F | df error |
|---|---|---|---|
| *R. biternatus* | 0.0001 | 1.55 | 125 |
| *R. pseudotrullifolius* | 0.021 | 1.28 | 88 |
| *P. antiscorbutica* | 0.0001 | 10.7 | 305 |

Table S2: ANOVA analyses indicating in *P. antiscorbutica* whether a) degree of phenotypic integration differs before and after application of treatments, b) phenotypic integration differs among treatments. P-values (P), F values (F) and error degree of freedom (df error) indicated.

| a) | P | F | df error |
|---|---|---|---|
| Warming | 0.58 | 0.58 | 9 |
| Shading | 0.13 | 1.7 | 9 |
| warming-shading | 0.4 | 0.89 | 9 |

| b) | P | F | dferror |
|---|---|---|---|
| Control vs warming | 0.84 | 0.04 | 18 |
| Control vs shading | 0.32 | 1.06 | 18 |
| Control vs warming-shading | 0.29 | 1.19 | 18 |
